# CDMAP/CDVIS: Context-dependent Mutation Analysis Package and Visualization Software

**DOI:** 10.1101/2022.02.03.479067

**Authors:** David L. Patton, Thomas Cardenas, Perrin Mele, Jon Navarro, Way Sung

## Abstract

The Context-dependent Mutation Analysis Package and Visualization Software (CDMAP/CDVIS) is an automated, modular toolkit used for analysis and visualization of context-dependent mutation patterns (site-specific variation in mutation rate from neighboring-nucleotide effects). The CDMAP algorithm calculates context-dependent mutation rates in chromosomes or replichores, and can generate high-resolution figures to analyze variation in mutation rate across spatiotemporal scales. Output from CDMAP can be integrated into CDVIS, an interactive database for visualizing mutation patterns across multiple taxa simultaneously.

## Introduction

Mutations are a primary source of genetic variation and understanding how, where, and when mutations arise is critical to elucidating the evolutionary process. Studying the rate and spectrum of spontaneous mutations can provide insight on how genomes evolve and adapt to changing environments. Spontaneous mutations are known to vary in size, scope, and type, with base substitutions and insertion-deletion mutations ranging from a single nucleotide to several thousand kilobases, and in some cases, entire chromosomes (Baer et al. 2007; Gordo et al. 2011; Heilbron et al. 2014; Keith et al. 2016; Wei et al. 2018; Sung et al. 2015).

Mutation accumulation (MA) studies, where organisms are bottlenecked to accumulate all but the most deleterious mutations (Lynch et al. 2016; Dillon et al. 2015), have provided a wealth of information regarding how organisms mutate. However, this data has also shown that mutation rate varies depending on genomic position (Foster et al. 2013), replication strand (Sung et al. 2015), mutation type (Long et al. 2014), and genomic context (Harris and Pritchard 2017; Schroeder et al. 2016; Long et al. 2014). Local-sequence context has been shown to influence site-specific mutation rates by up to 75-fold within the same sequence context (Dillon et al. 2015; Sung et al. 2015), and upwards of 403-fold within different contexts (Schroeder et al. 2016). Local sequence context has also been shown to have a large impact on site-specific mutation rates in bacteria, plants, and humans (Morton et al. 2006; Harris and Nielsen 2014; Harris 2015).

Although, evidence of context-specific mutation patterns has been observed across taxonomical life, our understanding of these patterns remains limited due to the ad-hoc methods employed in various studies that are designed specifically for a single organism (Lee et al. 2012; Dillon et al. 2018; Long et al. 2015b). Furthermore, these studies do not orient the mutations with respect to any genomic landmark (e.g., origin of replication) so it is nearly impossible to examine and contrast large-scale patterns driving spatiotemporal variation in mutation rate across multiple taxa.

To this extent, we have developed CDMAP, an analysis and visualization package to measure of the genome-wide rate and spectra of context-dependent mutations. CDMAP is a novel software package that can be used to categorizes mutations and their local-sequence context into a per-replichore or per-chromosome basis, generate estimates of context-dependent mutation rates, provide a graphical representation and statistical correlation of these rates to compare across multiple taxa, and the output data can be integrated into an interactive graphical database via CDVIS. CDMAP and CDVIS provide a uniform treatment for categorizing context-dependent mutation patterns that has not been previously established on an easily accessible platform within the open-source R environment. Being able to dissect mutational patterns via visualization tools can illuminate our understanding the mechanisms driving replication fidelity and genome evolution.

## Methods

The functionality of CDMAP and CDVIS are broken into three separate components of analysis. The first is the CDMAP Single Organism Analysis pipeline (SOA). The SOA pipeline provides the backbone analysis that catalogues nucleotide motifs across the genome, calculates context dependent mutation rates, and provides output csv files and organism-specific visualization output via Lattice (Sarkar 2008). The second component, CDMAP Multi Organism Analysis Pipeline (CDMAP-MOA), generates statistical correlations between different SOA analyses outlining potential relationships in contextual mutation patterns. The final component, CDVIS interfaces with CDMAP output to provide an accessible database of spatiotemporal variation in mutation patterns across analyzed genomes.

### Dependencies and Required Input Files

To identify context-dependent mutation patterns, we use the R-programming language, which is a robust library of bioinformatics, statistical, and data visualization packages. Several R dependencies are required for data preprocessing and postprocessing:

- SeqInR (OriLoc): R packages used to parse FASTA sequences and identify the origin of replication for strand specific analyses (Charif and Lobry 2007)
- Pracma: Numerical and statistical algorithms (Borchers 2021)
- Genbankr: Genbank file parsing (Lawrence 2019)
- Lattice: Lightweight data visualization package (Sarkar 2008)

Necessary input data for CDMAP includes a modified Variant Call File (VCF), the reference FASTA file, and an annotated GBK file. A VCF file is a space or tab-delimited file that can be generated by variant calling pipelines e.g., (SAMTOOLS/GATK) (Heng Li et al. 2009; Geraldine A. Van der Auwera 2020), containing the nucleotide position, the reference nucleotide, and the mutant nucleotide. The reference FASTA file is the genome sequence of the organism that corresponds to the nucleotide positions found in the VCF file. The annotated Genbank (GBK) file contains information about the location of genes in the reference FASTA. The reference FASTA, VCF, and GBK files used in the development of this package can be downloaded from the National Center for Biotechnology Information (NCBI) and the Short-read Archive (SRA) at NCBI.

### Replication Origin Determination and Replichore Partitioning

Context-specific mutation patterns have been shown to be asymmetrical with respect to the origin of replication (ORI) and replication terminus (TERM), such that the upstream 5’ and downstream 3’ base from the mutant site can influence site-specific mutation rate (Sung et al. 2015; Lee et al. 2012). CDMAP orients each mutation to a user-defined ORI location or an ORI defined by the OriLoc dependency (A. C. Frank 2000), an R package used to determine the minimum and maximum cumulative composite skew at synonymous sites (GC-skew) that is widely used to identify the ORI in bacterial organisms. CDMAP orients all variants with respect to the ORI for downstream analysis. After successful orientation and partitioning of the sequence data and mutations with respect to their ORI and TERM, nucleotide triplet counts (GWTC) for the chromosome and each replichore are tabulated for subsequent calculations.

### Nucleotide & Mutation Frequency Determination and Rate Analysis for Mutation Accumulation

To calculate the context-dependent mutation rate for mutation-accumulation experiments, genome-wide and replichore-wide triplet counts are counted (GWTC/RWTC). CDMAP then parses the variant call file to determine the upstream and downstream nucleotide associated with each variant and computes the mutation frequency and the context-dependent mutation rate at all 64 possible triplets:

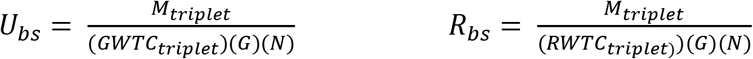

The context-dependent mutation rate for a triplet in the chromosome (*U*_*bs*_) is then determined by the by the total number of mutations observed at the center nucleotide of that triplet (*M*_*triplet*_) divided by the triplet count for the genome (*GWTC*_*triplet*_) the number of lineags (*N*) and the estimated number of generations elapsed (*G*). Replichore-specific rates are similarly calculated using mutations observed in a replichore (*R*_*bs*_) divided by the triplet count for that replichore (*RWTC*_*triplet*_), *G*, and *N*. In addition to a triplet frame, CDMAP accounts for and tracks data regarding upstream and downstream neighboring nucleotides in a 5-mer reference frame, i.e., NXNY downstream and YNXN nucleotide upstream contexts, where X and Y are mutable nucleotides, and N represents a neighbor nucleotide of X.

### Multi-Organism Analysis

CDMAP was developed to allow for flexibility in the number of organisms analyzed. During the run process for a single organism, CDMAP dynamically creates a repository of the output which can be used for downstream comparison against additional CDMAP runs. Once selected genomes have been analyzed, the user can perform a Multi-Organism Analysis to compare the context-dependent mutation patterns generated using the SOA pipeline. This comparison can occur on a chromosome-wide, strand-specific, or replichore-specific basis. CDMAP performs a one-to-many Pearson’s correlation sequentially with each organism, and automatically orients the coefficients according to GC content for display as heat maps in lattice.

### CDMAP-SOA Visualization

Complex patterns within large-scale data sets are often easier to identify using visualization tools. Relevant information about triplet frequency, variant distribution, and genome-wide and replichore-specific mutation rates are passed through Lattice and correlation between input files can be automatically formatted (Fig. 1A) (Sarkar 2008). Throughout the process, CDMAP collects and outputs both CSV format spreadsheets and heatmaps in dynamically generated output directories that are categorized for easy navigation and downstream analyses.

**Figure 1:**
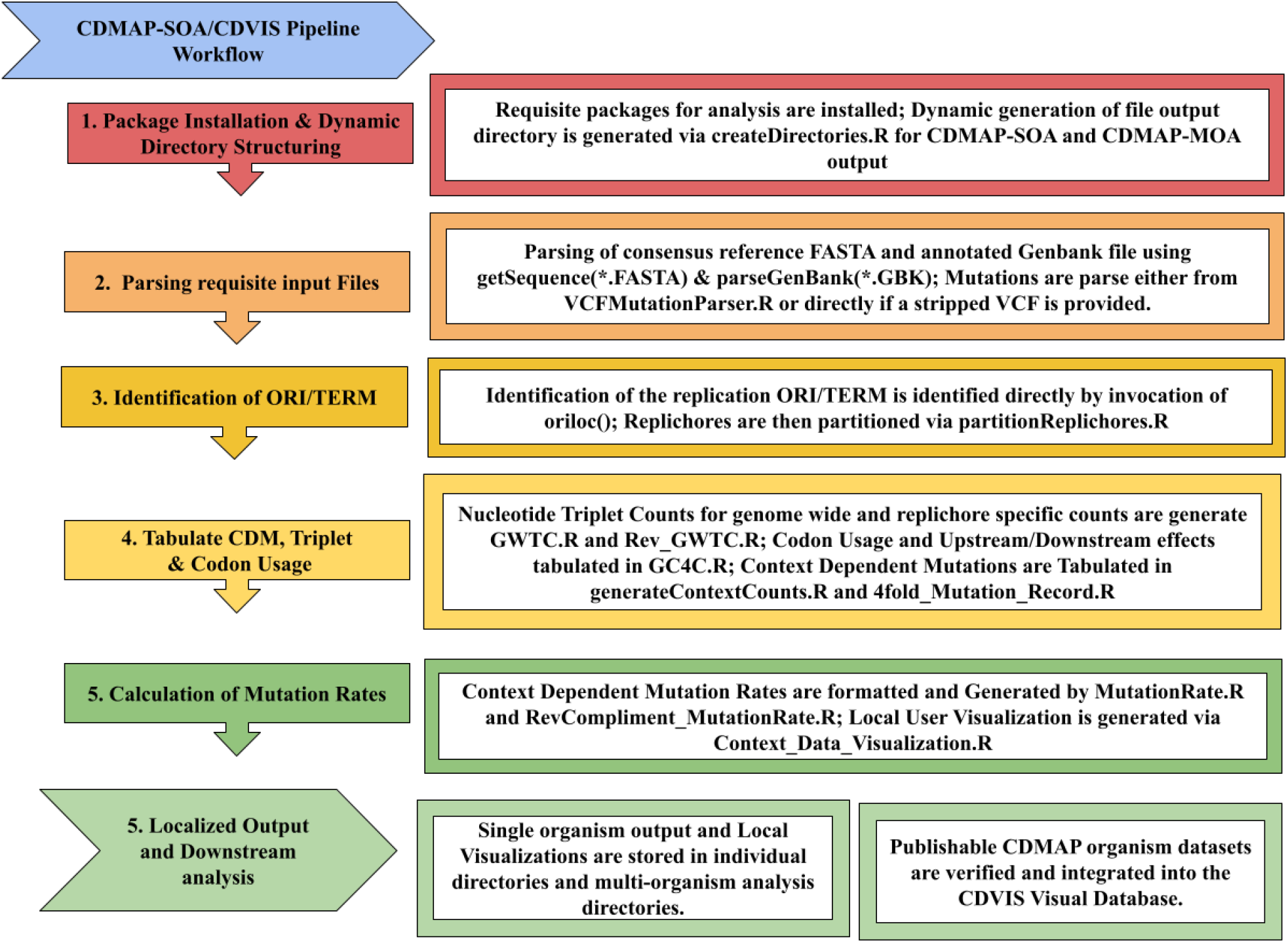
CDMAP/CDVIS Diagram Workflow – Visual outlining the major steps taken during the analysis of the CDMAP-SOA pipeline for generation of context dependent mutation rates, genome-wide triplet and codon usage counts on a per-chromosome or per replichore basis.

In the example shown in Figure 2, CDMAP has generated the context-dependent mutation rates for all 64 nucleotide triplets from a mismatch repair deficient line of *B. subtilis* (Sung et al. 2015). In this example site-specific rates are shown for the left and right replichore, with each context oriented so that both strands are synthesized identically (mutations and contexts are taken with respect to their reverse complement). Our algorithm generates identical context-dependent mutation rates when compared to ad-hoc methods used in previous studies (Sung et al. 2015).

**Figure 2.**
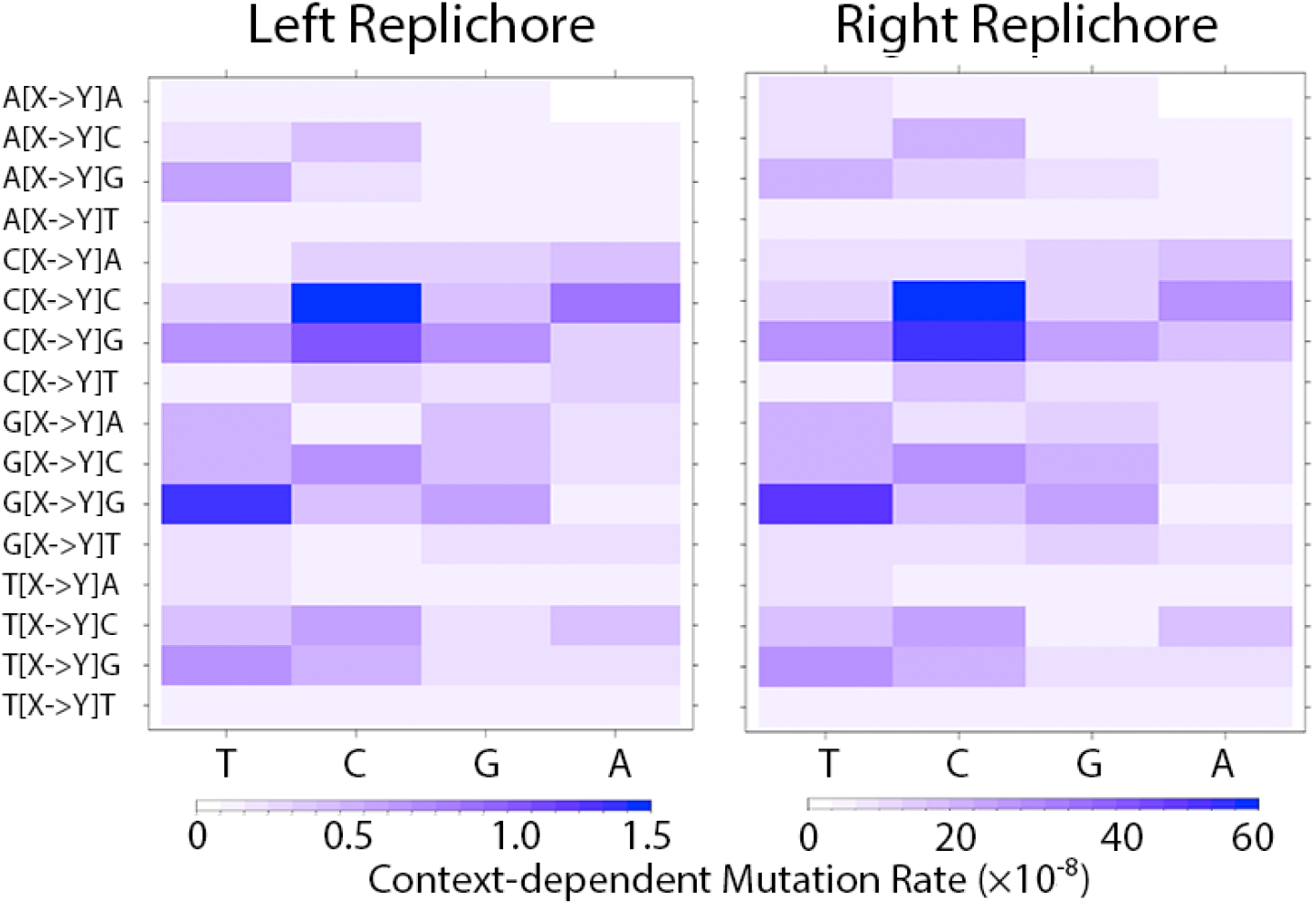
CDMAP Single Organism Analysis (SOA) output of *Bacillus subtilis* mismatch repair deficient MA lines. Context-dependent mutation rates shown for left replichore and right replichore. Each row repesents a triplet N[X->Y]N with N as the left and right neighboring nucleotide and X and Y repesenting the reference nucleotide and mutation respectively.

### CDMAP-MOA Visualization

In figure 3A, we show an example of multiple organisms benchmarked via CDMAP-SOA (Table S1). In this visualization, columns are ranked from AT rich (top) to GC-rich (bottom) and rows are oriented GC rich (left) to AT-rich (right). The one-to-one comparison context dependent mutation rates between organisms are color coordinated relative to its Pearson’s coefficient, as indicated by the heat map legend (Figure 3A).

**Figure 3.**
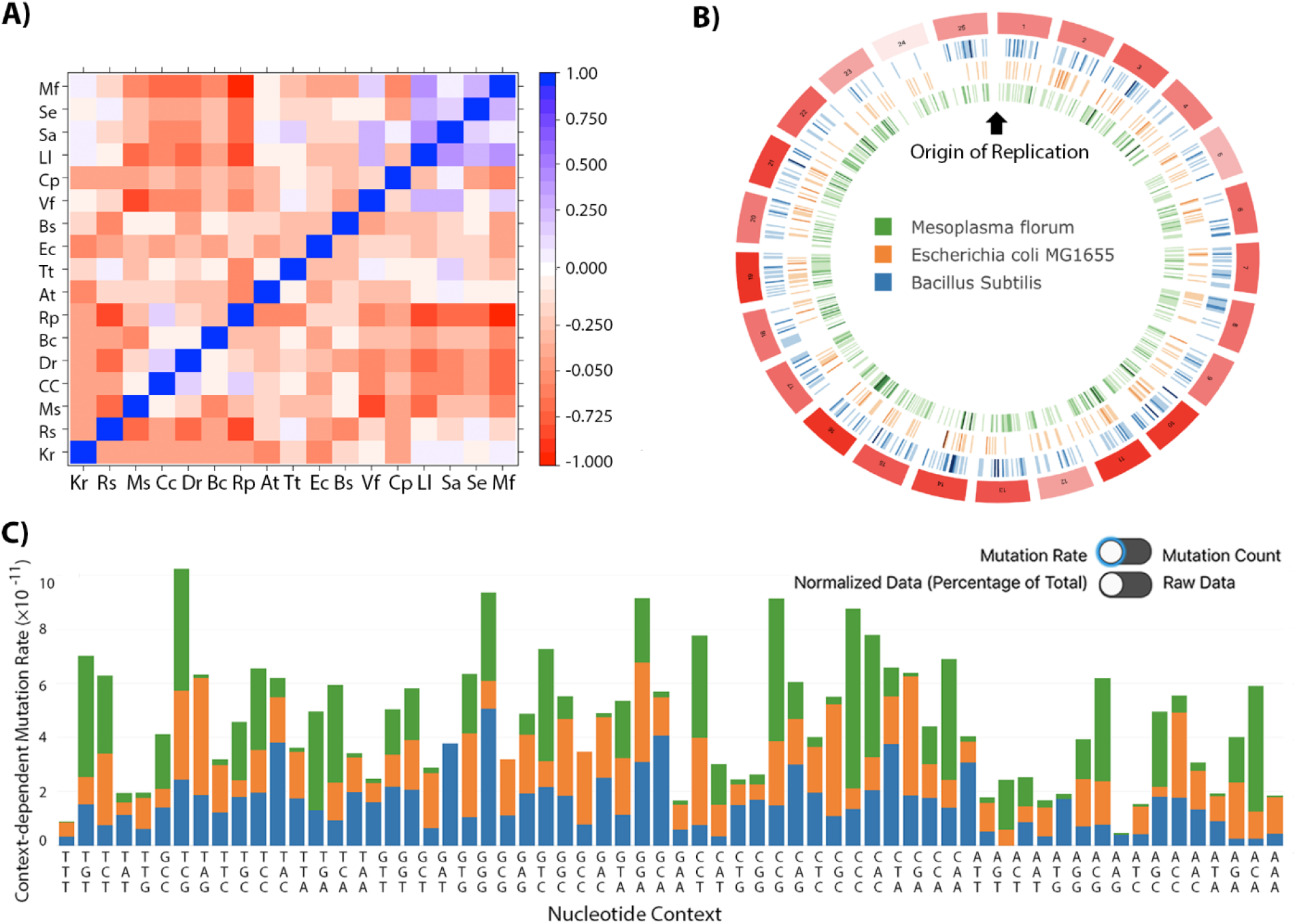
CDMAP / CDVIS Webtool: A) CDMAP One-to-Many Correlation Heatmap for the primary chromosome of 17 bacterial mutation accumulation datasets (Pearson’s correlation coefficient sorted by coding region GC content – Table S1). B) CDVIS online CIRCOS (Krzywinski et al. 2009) visualization of *B. subtilis, E. coli, and M. florum*. Mutations for each organism are oriented into 25 bins with the origin of replication located at bin 1 and the terminus located in bin 13/14. Intensity of tick marks show the number of mutations for that species, and intensity of outside red boxes indicate increasing density of mutations across selected organisms. Visualization tools can display cumulative mutation data across multiple organisms for each selected bin (www.wsunglab.com:3000). C) 64 codon triplet stacked graph indicating the context-dependent mutation rate of each wild-type organism and can be displayed as a rate, raw counts, or conditional rate per triplet.

### Web Visualization Using CDVIS

Data generated through the CDMAP pipeline can integrated into our front-facing web server, the Context-dependent Visualization Software (CDVIS). CDVIS contains CDMAP output from existing MA experiments that can be used as a comparative-framework against future datasets. CDVIS takes output from the CDMAP pipeline and organizes them into JSON objects that are then are dynamically loaded into a circular format using CIRCOS (Martin Krzywinski 2009) (Fig. 3B).

The user can select numerous datasets from available pre-loaded MA experiments to visualize the spatiotemporal variation of mutation rate in MA organisms (www.wsunglab.com:3000). Each circular track represents a single organism, and tick marks within the tracks indicate the location and density of mutations at that location. The genome is divided into selectable bins (size 25/50/75) and the density/type/and rate of mutations of the selected organism(s) are displayed in a side panel. For easy comparison, CDMAP/CDVIS displays the first bin starting at the origin of replication. Finally, the visualization tools allow for the summation of conditional mutation rates (mutation rates normalized to genome-wide nucleotide content - Fig. 3B) and cumulative context-dependent mutation rates for different triplets (Fig. 3C). At this time, additions to the available visualized organisms can be made through email requests to the authors.

## Results and Discussion

CDMAP was quantitatively benchmarked against 17 mutation accumulation datasets in prokaryotic organisms (Sung et al. 2015; Lynch et al. 2016; Long et al. 2014; Kucukyildirim et al. 2016b; Dillon et al. 2015; Sung et al. 2012a; Gilchrist et al. 2015; Lynch 1996), which harbor a variety of different genomic architectural features (Table S1). The majority of MA studies contain organisms with a singular, circular chromosome such as *Escherichia coli*, while others may have multiple genomic elements such as chromids and plasmids (Table S1), or may be deficient in repair enzymes such as mismatch repair. We processed 17 organisms (Table S1) containing a total of 12493 mutations. There are no other software toolkits that are readily available for comparison, but we were able to recover identical context-dependent mutation rates from prior studies generated using ad-hoc methods (Sung et al. 2015; Lee et al 2012). All data was uploaded to the CDVIS visualization tool at www.wsunglab.com:3000.

CDMAP was designed to be a lightweight analysis package capable of running on a standard laptop or desktop. Each of the 17 datasets were analyzed on an iMac with a 2.9Ghz quad core intel i5 processor, 16GB 1600Mhz DDR3 ram, running MAC OSX Catalina. On the benchmarked machine, CDMAP utilized approximately 1gb (6.25% memory, 7 threads) and roughly 60% CPU utilization during its most computationally intensive processes. The average runtime of a given organism came in around 90 minutes for an average size bacterial genome (~5mb).

### Practical Example

The following commands can be used as a practical example on how CDMAP can be used to generate and analyze context-dependent mutation patterns in genomic data. In this short tutorial we will walk through basic commands used to generate context-dependent mutation patterns from a *Bacillus subtilis* mutation accumulation dataset *(Sung et al. 2015)*. This example data for *B. subtilis* and other organisms used for benchmarking are included with CDMAP package found on the Github repository found at (https://github.com/DLP-Informatics/CDMAP).

To begin running CDMAP, first install R (https://www.r-project.org/), then navigate to the directory in which you unpacked the CDMAP package and execute the following command in terminal:

> *>Rscript CDMAP_SingleOrganismAnalysis.R*

The user will be prompted for the name of the output folder designated by the end user, whether the user is using a VCF or modified base call file, and the full path location of the reference sequence, genbank file, and the VCF or modified base call file.

> *>Bacillus_subtilis_WT*
>
> *>basecall*
>
> >/Users/Username/Desktop/CDMAP/Test_Datasets/bacillus/Bacillus_3610.fasta
>
> >/Users/Username/Desktop/CDMAP/Test_Datasets/bacillus/NC000964.gbk
>
> >/Users/Username/Desktop/CDMAP/Test_Datasets/bacillus/Bacillus_WT.csv

CDMAP will then prompt the user for how many generations have elapsed and how many lineages were in the experiment. For analyzing data that is not from a mutation accumulation experiment, generations can be estimated using a molecular clock method. By default, mutation rates will be scaled to 1×10^−8^ for ease of visualization in lattice, but scaling can be changed by the user (0 for default, 1 for scaling to the average mean of rates, or 2 for custom parameters). Finally, the user can allow for the automatic calculation of the location of the ORI using ORIloc (auto) or manually input the location of the ORI. In our example, the *B. subtilis* MA experiment underwent 5077 generations across 50 lines).

> >*5077*
>
> >*50*
>
> >0
>
> >auto

After these following steps have been completed, CDMAP-SOA will have the requisite information needed to count contexts, and automatically estimate mutation rates for the chromosome, for each replichore (on both strands), and generate high-resolution heatmaps that can be accessed by lattice (Fig. 2). Once the run is complete, all SOA output files will be placed into the CDMAP_Output/Output_Directory that was designated by the user. Upon request, thits data can be interfaced into CDVIS for further analysis. If the user wants to perform direct correlation between the rates from different organisms or experiments, they can invoke CDMAP-MOA using the following command:

> *Rscript CDMAP_MultiOrganism_Analysis.R*

CDMAP-MOA will correlate analyze all SOA runs located within the specified Output_Directory, sort the organisms by GC-content, and generate a high-resolution correlation heatmap for downstream analysis (Fig. 3A). An in-depth description of the files generated, information on the directory structure, and a full technical document can be found in the CDMAP technical document which can be found in the package as well as at the github repository (https://github.com/DLP-Informatics/CDMAP/documentation).

## Conclusion

CDMAP is a toolkit designed to streamline the analysis of context-dependent mutations from genomic sequence data. While CDMAP has been benchmarked on bacterial mutation accumulation datasets with a single replication origin, CDMAP is capable of analyzing linear chromosomes, including viral, archaeal, and eukaryotic datasets with the caveat of manually inputting the ORI. Determining the ORI in non-prokaryotic chromosomes can be done in a few different ways, including replication profile construction via deep sequencing methods (Jia Xu 2012). In addition, CDMAP can not only be applied to mutation datasets but also to silent sites from population sequencing. Application of CDMAP on data from natural populations and integration into CDVIS can assist researchers in exploring how spatiotemporal variation in mutation rate can drive genome evolution.

## Data Availability

The CDMAP source code is freely available for non-commercial academic use at https://github.com/DLP-Informatics/CDMAP; data is also available at CDVIS for viewing at: https://www.wsunglab.com:3000

---

**Supplemental Table 1.** Mutation accumulation datasets used during the development of CDMAP, and generated visualization data can be accessed using CDVIS (www.wsunglab.com:3000). Table headers are species name, accession GCF or GCA number of the reference genome assembly used in the analysis (FASTA/GBK files), # of MA generations, # of MA lineages, chromosomes, total observed mutations, GC-content and the kilo base location of the origin of replication (ORI). MMR-indicates mismatch-repair deficient strain(Senra et al. 2018; Dillon et al. 2017; Sun et al. 2017; Kucukyildirim et al. 2016a; Sung et al. 2016; Kucukyildirim et al. 2016b; Lee et al. 2016; Sung et al. 2015; Long et al. 2015a; Sung et al. 2012b; Long et al. 2018; Dillon et al. 2015).

| Species | Acc #. GCF/GCA: | Gen. | Lines | Chr. | Mut. | GC% | ORI (KB) |
| --- | --- | --- | --- | --- | --- | --- | --- |
| *Agrobacterium tumefaciens* C58 | 000092025.1_ASM9202v1 | 5819 | 47 | 2 | 233 | 60 | 2765 |
| *Bacillus subtilis* NCIB 3610 | 000186085.1_ASM18608v1 | 5077 | 50 | 1 | 350 | 44 | 4170 |
| *Bacillus subtilis* NCIB 3610 (MMR-) | 000186085.1_ASM18608v1 | 5077 | 50 | 1 | 5295 | 44 | 4170 |
| *Burkholderia cenocepacia* HI2424 | 000203955.1_ASM20395v1 | 5554 | 47 | 3 | 130 | 67 | 69 |
| *Caulobacter crescentus* NA1000 | 000022005.1_ASM2200v1 | 4284 | 44 | 1 | 259 | 68 | 3818 |
| *Colwellia psychrerythraea* 34H | 000012325.1_ASM1232v1 | 1078 | 84 | 1 | 400 | 39 | 4903 |
| *Deinococcus radiodurans* BAA-816 | 001638825.1_ASM163882v1 | 5961 | 43 | 2 | 331 | 68 | 258 |
| *Escherichia coli* K-12 MG1655 | 000005845.2_ASM584v2 | 1682 | 46 | 1 | 1623 | 52 | 3644 |
| *Escherichia coli* K-12 MG1655 (MMR -) | 000005845.2_ASM584v2 | 1682 | 46 | 1 | 231 | 52 | 3644 |
| *Kineococcus radiotolerans* SRS30216 | 000017305.1_ASM1730v1 | 4724 | 44 | 1 | 280 | 74 | 522 |
| *Lactococcus lactis* DSMZ20481 | 900088425.1_A12 | 3973 | 63 | 1 | 813 | 37 | 1 |
| *Mesoplasma florum* L1 | 000479355.1_ASM47935v1 | 2351 | 28 | 1 | 544 | 27 | 326 |
| *Mycobacterium smegmatis* MC2 155 | 000283295.1_ASM28329v1 | 4900 | 49 | 1 | 856 | 68 | 3609 |
| *Rhodobacter sphaeroides* ATCC 17025 | 000016405.1_ASM1640v1 | 4544 | 46 | 1 | 107 | 69 | 2 |
| *Ruegeria pomeryoi* DSS-3 | 000011965.2_ASM1196v2 | 5386 | 47 | 1 | 147 | 65 | 1 |
| *Staphylococcus aureus* ATCC 25923 | 000756205.1_ASM75620v1 | 2716 | 83 | 1 | 274 | 34 | 3 |
| *Staphylococcus epidermis* ATCC 122228 | 000007645.1_ASM764v1 | 7101 | 22 | 1 | 294 | 33 | 2419 |
| *Teredinibacter turnerae* T7901 | 000023025.1_ASM2302v1 | 3025 | 42 | 1 | 779 | 52 | 4252 |
| *Vibrio fischeri* ES114 | 000011805.1_ASM1180v1 | 5187 | 48 | 2 | 132 | 40 | 1 |
| *Vibrio fischeri* ES114 (MMR -) | 000011805.1_ASM1180v1 | 5187 | 48 | 2 | 2909 | 40 | 1 |

## Literature Cited

A. C. Frank, J.R.L., 2000 Oriloc: prediction of replication boundaries in unannotated bacterial chromosomes. Bioinformatics 16 (6):560–561.

Baer, C.F., M.M. Miyamoto, and D.R. Denver, 2007 Mutation rate variation in multicellular eukaryotes: causes and consequences. Nat Rev Genet 8 (8):619–631.

Borchers, H.W., 2021 pracma: Practical Numerical Math Functions.

Charif, D., and J.R. Lobry, 2007 SeqinR 1.0-2: A Contributed Package to the R Project for Statistical Computing Devoted to Biological Sequences Retrieval and Analysis, pp. 207–232 in Structural Approaches to Sequence Evolution: Molecules, Networks, Populations, edited by U. Bastolla, M. Porto, H.E. Roman and M. Vendruscolo. Springer Berlin Heidelberg, Berlin, Heidelberg.

Dillon, M.M., W. Sung, M. Lynch, and V.S. Cooper, 2015 The rate and molecular spectrum of spontaneous mutations in the GC-rich multichromosome genome of Burkholderia cenocepacia. Genetics 200 (3):935–946.

Dillon, M.M., W. Sung, M. Lynch, and V.S. Cooper, 2018 Periodic Variation of Mutation Rates in Bacterial Genomes Associated with Replication Timing. MBio 9 (4).

Dillon, M.M., W. Sung, R. Sebra, M. Lynch, and V.S. Cooper, 2017 Genome-Wide Biases in the Rate and Molecular Spectrum of Spontaneous Mutations in Vibrio cholerae and Vibrio fischeri. Mol Biol Evol 34 (1):93–109.

Foster, P.L., A.J. Hanson, H. Lee, E.M. Popodi, and H. Tang, 2013 On the mutational topology of the bacterial genome. G3 (Bethesda) 3 (3):399–407.

Geraldine A. Van der Auwera, B.D.O.C., 2020 Genomics in the Cloud: Using Docker, Gatk and WDL in Terra.

Gilchrist, M.A., W.C. Chen, P. Shah, C.L. Landerer, and R. Zaretzki, 2015 Estimating Gene Expression and Codon-Specific Translational Efficiencies, Mutation Biases, and Selection Coefficients from Genomic Data Alone. Genome Biol Evol 7 (6):1559–1579.

Gordo, I., L. Perfeito, and A. Sousa, 2011 Fitness Effects of Mutations in Bacteria. Journal of Molecular Microbiology and Biotechnology 21 (1-2):20–35.

Harris, K., 2015 Evidence for recent, population-specific evolution of the human mutation rate. Proc Natl Acad Sci U S A 112 (11):3439–3444.

Harris, K., and R. Nielsen, 2014 Error-prone polymerase activity causes multinucleotide mutations in humans. Genome Res 24 (9):1445–1454.

Harris, K., and J.K. Pritchard, 2017 Rapid evolution of the human mutation spectrum. Elife 6.

Heilbron, K., M. Toll-Riera, M. Kojadinovic, and R.C. MacLean, 2014 Fitness is strongly influenced by rare mutations of large effect in a microbial mutation accumulation experiment. Genetics 197 (3):981–990.

Heng Li, B.H., Alec Wysoker, Tim Fennell, Jue Ruan, Nils Homer, G.A. Gabor Marth, Richard Durbin,∗ and 1000 Genome Project Data, and P. Subgroup, 2009 The Sequence Alignment/Map format and SAMtools. Bioinformatics 25 (16):2078–2079.

Jia Xu, Y.Y., Alexander M Tsankov, et al., 2012 Genome-Wide Identification and characterization of replication origins by deep sequencing. Genome Biology 13 (April 2012):14.

Keith, N., A.E. Tucker, C.E. Jackson, W. Sung, J.I. Lucas Lledo et al., 2016 High mutational rates of large-scale duplication and deletion in Daphnia pulex. Genome Res 26 (1):60–69.

Krzywinski, M., J. Schein, I. Birol, J. Connors, R. Gascoyne et al., 2009 Circos: an information aesthetic for comparative genomics. Genome Res 19 (9):1639–1645.

Kucukyildirim, S., H. Long, W. Sung, S.F. Miller, T.G. Doak et al., 2016a The Rate and Spectrum of Spontaneous Mutations in Mycobacterium smegmatis, a Bacterium Naturally Devoid of the Post-replicative Mismatch Repair Pathway. G3 (Bethesda).

Kucukyildirim, S., H. Long, W. Sung, S.F. Miller, T.G. Doak et al., 2016b The Rate and Spectrum of Spontaneous Mutations in Mycobacterium smegmatis, a Bacterium Naturally Devoid of the Postreplicative Mismatch Repair Pathway. G3 (Bethesda) 6 (7):2157–2163.

Lawrence, G.B.a.M., 2019 genbankr: Parsing GenBank files into semantically useful objects. R package version 1.14.0.

Lee, H., T.G. Doak, E. Popodi, P.L. Foster, and H. Tang, 2016 Insertion sequence-caused large-scale rearrangements in the genome of Escherichia coli. Nucleic Acids Res 44 (15):7109–7119.

Lee, H., E. Popodi, H. Tang, and P.L. Foster, 2012 Rate and molecular spectrum of spontaneous mutations in the bacterium Escherichia coli as determined by whole-genome sequencing. Proc Natl Acad Sci USA 109 (41):e2774–e2783.

Long, H., S. Kucukyildirim, W. Sung, E. Williams, H. Lee et al., 2015a Background Mutational Features of the Radiation-Resistant Bacterium Deinococcus radiodurans. Mol Biol Evol 32 (9):2383–2392.

Long, H., W. Sung, S. Kucukyildirim, E. Williams, S.F. Miller et al., 2018 Evolutionary determinants of genome-wide nucleotide composition. Nat Ecol Evol 2 (2):237–240.

Long, H., W. Sung, S.F. Miller, M.S. Ackerman, T.G. Doak et al., 2014 Mutation rate, spectrum, topology, and context-dependency in the DNA mismatch repair-deficient Pseudomonas fluorescens ATCC948. Genome Biol Evol 7 (1):262–271.

Long, H., W. Sung, S.F. Miller, M.S. Ackerman, T.G. Doak et al., 2015b Mutation rate, spectrum, topology, and context-dependency in the DNA mismatch repair-deficient Pseudomonas fluorescens ATCC948. Genome Biol Evol 7 (1):262–271.

Lynch, M., M.S. Ackerman, J.F. Gout, H. Long, W. Sung et al., 2016 Genetic drift, selection and the evolution of the mutation rate. Nat Rev Genet 17 (11):704–714.

Lynch, T.T.K.M., 1996 Estimate of the genomic mutation rate deleterious to overall fitness in E. coli. Nature 381 (20):694–696.

Martin Krzywinski, J.S., Inancx Birol, Joseph Connors, Randy Gascoyne, Doug Horsman, Steven J. Jones, and Marco A. Marra, 2009 Circos: An information aesthetic for comparative genomics. Genome Research.

Morton, B.R., I.V. Bi, M.D. McMullen, and B.S. Gaut, 2006 Variation in mutation dynamics across the maize genome as a function of regional and flanking base composition. Genetics 172 (1):569–577.

Sarkar, D., 2008 Lattice: Multivariate Data Visualization with R: Springer.

Schroeder, J.W., W.G. Hirst, G.A. Szewczyk, and L.A. Simmons, 2016 The Effect of Local Sequence Context on Mutational Bias of Genes Encoded on the Leading and Lagging Strands. Curr Biol 26 (5):692–697.

Senra, M.V.X., W. Sung, M. Ackerman, S.F. Miller, M. Lynch et al., 2018 An Unbiased Genome-Wide View of the Mutation Rate and Spectrum of the Endosymbiotic Bacterium Teredinibacter turnerae. Genome Biol Evol 10 (3):723–730.

Sun, Y., K.E. Powell, W. Sung, M. Lynch, M.A. Moran et al., 2017 Spontaneous mutations of a model heterotrophic marine bacterium. The ISME journal 11 (7):1713–1718.

Sung, W., M.S. Ackerman, M.M. Dillon, T.G. Platt, C. Fuqua et al., 2016 Evolution of the Insertion-Deletion Mutation Rate Across the Tree of Life. G3 (Bethesda) 6 (8):2583–2591.

Sung, W., M.S. Ackerman, J.F. Gout, S.F. Miller, E. Williams et al., 2015 Asymmetric context-dependent mutation patterns revealed through mutation-accumulation experiments. Mol Biol Evol 32 (7):1672–1683.

Sung, W., M.S. Ackerman, S.F. Miller, T.G. Doak, and M. Lynch, 2012a Drift-barrier hypothesis and mutation-rate evolution. Proc Natl Acad Sci U S A 109 (45):18488–18492.

Sung, W., M.S. Ackerman, S.F. Miller, T.G. Doak, and M. Lynch, 2012b Drift-barrier hypothesis and mutation-rate evolution. Proc Natl Acad Sci USA 109 (45):18488–18492.

Wei, W., L. Xiong, Y.N. Ye, M.Z. Du, Y.Z. Gao et al., 2018 Mutation Landscape of Base Substitutions, Duplications, and Deletions in the Representative Current Cholera Pandemic Strain. Genome Biol Evol 10 (8):2072–2085.

